# Silencing NADPH-Cytochrome P450 reductase affects imidacloprid susceptibility, fecundity, and embryonic development in *Leptinotarsa decemlineata*

**DOI:** 10.1101/2020.09.29.318634

**Authors:** Timothy W. Moural, Liping Ban, Jonathan A. Hernandez, Meixiang Wu, Chaoyang Zhao, Subba R. Palli, Andrei Alyokhin, Fang Zhu

**Author notes:** These authors contributed equally to this work (Timothy W. Moural, Liping Ban). Corresponding author. *Email address:* (F. Zhu).

## Abstract

The Colorado potato beetle (CPB) is a prominent insect pest of potatoes, tomatoes and eggplants all over the world, however, the management of CPB remains a challenging task for more than one hundred years. We have successfully developed bacteria-expressed dsRNA-mediated feeding RNA interference (RNAi) approach in our previous study. A critical step towards field management of CPB via feeding RNAi is to identify effective and environmentally safe target genes. NADPH-Cytochrome P450 reductase (CPR) plays a central role in cytochrome P450 action. The full length *Leptinotarsa decemlineata* CPR (*LdCPR*) cDNA was isolated from an imidacloprid resistant population. The *LdCPR* gene was ubiquitously expressed in all stages tested but showed an increase in expression during the early stage of embryonic development. The bacteria-expressed dsRNA-mediated feeding RNAi of *LdCPR* in adults caused systemic knock down expression of the gene coding for *LdCPR* in both adults and their eggs. Suppression of *LdCPR* expression increased susceptibility of imidacloprid in resistant beetles, as well as a significant decrease of fecundity in female beetles (29% less eggs/day) and the hatching rate (47%) of their eggs. These data suggest that *LdCPR* plays important roles in insecticide detoxification and biosynthetic pathways of endogenous compounds and may serve as an essential target to control CPB.

**HIGHLIGHTS:**

- High expression of *LdCPR* was observed in the egg stage.
- Silencing of *LdCPR* reduced the CPR enzymatic activities.
- *LdCPR* knockdown increased imidacloprid susceptibility.
- *LdCPR* knockdown decreased the fecundity and enhanced embryonic lethality.

## 1. Introduction

Cytochrome P450s constitute one of the largest and oldest superfamilies of enzymes that catalyze extraordinarily diverse chemical reactions, from hydroxylation to epoxidation, O-, N-, and S-dealkyation, N- and S-oxidations, and up to as many as 60 (Coon et al., 1996; Feyereisen, 1999). Due to the remarkably flexible substrate specificity and catalytic versatility, cytochrome P450s play a central role for insects in adaptation to numerous chemical stresses from natural and anthropogenic xenobiotics (Krieger et al., 1971; Feyereisen et al., 1989; Cariño et al., 1992; Schuler, 1996; David et al., 2013; Zhu et al., 2014). Insect P450s are also involved in the biosynthesis and/or degradation of endogenous compounds, including hormones (Helvig et al., 2004a; Rewitz et al., 2007), fatty acids (Helvig et al., 2004b), cuticle hydrocarbon and other pheromones (Maibeche-Coisne et al., 2004; Sandstrom et al., 2006; Qiu et al., 2012; Balabanidou et al., 2016), which are extremely critical for insects to cope with their environment.

To complete the catalytic cycle, cytochrome P450s require the delivery of two electrons (Feyereisen, 2012). In the endoplasmic reticulum, the electrons for this process are transferred from NADPH to the P450 heme co-factor by an obligated electron transporter, NADPH-Cytochrome P450 reductase (CPR) (Feyereisen, 2012). In certain situations, an additional potential electron donor, microsomal cytochrome b5 may increase the transfer rate of the second electron (Feyereisen, 2012; Waskell and Kim, 2015). CPR belongs to a family of flavoproteins that use flavin mononucleotide (FMN) and flavin adenine dinucleotide (FAD) as cofactors. Based on X-ray crystallography of the rat CPR (Wang et al., 1997), insect CPRs are composed of three major domains, the FMN, FAD and NADP(H) binding domains (Feyereisen, 2012). Studies suggested that *CPR* genes evolved from the fusion of two ancestral genes: one coding for a ferredoxin reductase with FAD and NADP(H) domains and the other coding for a flavodoxin with FMN domain (Porter and Kasper, 1986). CPR serves as a redox partner for microsomal cytochrome P450s and several other microsomal oxygenase enzymes in most eukaryotic cells. These enzymes include but are not limited to cytochrome b5, squalene monooxygenase, and heme oxygenase (Feyereisen, 1999; Riddick et al., 2013).

Each insect genome contains about one hundred microsomal cytochrome P450 genes but generally only one CPR gene (Zhu et al., 2012). In insect microsomes, both P450 and CPR enzymes are integral membrane proteins. The ratio of these two enzymes is about 6-18 to 1 in insects (Feyereisen, 2005). As inhibition or silence of CPR inactivates all microsomal P450s (Henderson et al., 2003), it offers a unique way to decipher contributions of cytochrome P450s in xenobiotic adaptation and/or endogenous compounds biosynthesis or degradation. For example, silencing CPR in *Anopheles gambiae* and *Cimex lectularius* increased insect susceptibilities to pyrethroid insecticides, suggesting a key role that cytochrome P450s play in pyrethroid metabolism (Lycett et al., 2006; Zhu et al., 2012). In *Bactrocera dorsalis, Nilaparvata lugens, Rhopalosiphum padi*, *Laodelphax striatellus*, *Locusta migratoria*, and *Aphis citricidus*, knockdown of CPR resulted in enhanced susceptibilities to malathion, beta-cypermethrin and imidacloprid, isoprocarb and imidacloprid, buprofezin, carbaryl, and abamectin, respectively (Huang et al., 2015; Liu et al., 2015; Wang et al., 2016; Zhang et al., 2016; Zhang et al., 2017; Jing et al., 2018). These studies revealed the involvement of cytochrome P450 mediated detoxification in insecticide adaptation. In addition, *CPR*-silenced *Drosophila melanogaster* showed a deficiency in cuticular hydrocarbons and reduced viability upon adult emergence, suggesting the involvement of cytochrome P450s in cuticular hydrocarbon biosynthesis (Qiu et al., 2012).

The Colorado potato beetle (CPB), *Leptinotarsa decemlineata*, is a destructive phytophagous pest causing serious defoliation of potatoes, tomatoes, and eggplants all over the world (Schoville et al., 2018). For a long time, management of CPB has remained a difficult task due to many factors associated with this beetle, such as the high fecundity, flexible life history, and remarkable capability to adapt to numerous biotic and abiotic stresses (Alyokhin et al., 2008; Alyokhin et al., 2015; Ma et al., 2019). In recent years, advanced technologies in genome sequencing and functional genomics have facilitated multiple feasible approaches for CPB control, including bacteria-expressed double stranded RNA-mediated feeding RNA interference (RNAi) (Zhu et al., 2011; Palli, 2014; Zhu et al., 2014; San Miguel and Scott, 2016) and genetic modification of potato plants (Zhang et al., 2015). However, one of the critical steps towards successful field practice is to identify effective and environmentally safe target genes for discovery of novel RNAi-based biopesticides (Lundgren and Duan, 2013; Zhu et al., 2014; Ma et al., 2019; Zhu and Palli, 2020).

In order to effectively control this pest, we have successfully developed a bacteria-expressed double stranded RNA-mediated feeding RNA interference (RNAi) approach, which works in a target-specific, highly efficacious, and cost-effective manner for CPB control (Zhu et al., 2011; Kadoić Balaško et al., 2020). Resistance mechanism studies revealed that the cytochrome P450 monooxygenase-mediated detoxification is the most common mechanism of resistance across many populations from diverse geographical origins (Alyokhin et al., 2008; Clements et al., 2016; Zhu et al., 2016). With critical biological function associated with cytochrome P450s and other oxygenase enzymes, CPR will be an essential target for the development of novel RNAi-based biopesticides towards effective CPB management (Zhu et al., 2012; Huang et al., 2015; Shi et al., 2015; Zhang et al., 2017). In current study, the full-length cDNA of *Leptinotarsa decemlineata* CPR (*LdCPR*) was isolated and cloned from an imidacloprid resistant population. The developmental expression of the *LdCPR* gene in all life stages was investigated. The effects and mechanisms of bacteria-expressed dsRNA-mediated feeding RNAi of *LdCPR* in adults and the parental RNAi effects in CPB eggs were also examined. The possibilities for developing *LdCPR* as an essential target to control CPB are discussed.

## 2. Materials and methods

### 2.1. Insects

The susceptible CPB was kindly provided by Dr. Don Weber of USDA-ARS, originally supplied by the New Jersey Department of Agriculture. The imidaclorpid resistant CPB population was collected from Long Island, NY (Zhu et al., 2016). Both populations were reared on Red Norland potato plants in several BugDorm insect cages (MegaView Science Education Services Co., Ltd.) at 25±5 °C under a light:dark regimen of 16:8 h in the greenhouse. New potted potato plants were provided for each cage twice every week. Egg masses were collected each day and stored in a petri dish with water-soaked filter paper and kept in an incubator (25±1 °C, RH of 70%, L:D=16:8). After hatching, the young larvae were fed fresh potato leaves. The 2^nd^ instar larvae were transferred back to the green house.

### 2.2. RNA extraction, cDNA synthesis, and LdCPR cloning

Total RNA was isolated from 3 to 30 CPB beetles in each life stage (1 day and 5 day old eggs, 1^st^-4^th^ instar larvae, pupae, 1-week old female and male adults) with the TRIzol reagent (Invitrogen) and treated with DNase I (Invitrogen) according to the manufacturer’s protocols. The quality and quantity of total RNA were determined with a NanoDrop One Microvolume UV-Vis Spectrophotometer (Thermo Fisher). cDNA was synthesized using iScript cDNA synthesis kit (Bio-Rad Laboratories, Hercules, CA) following the protocol of manufacturer. A cDNA mixture of all life stages was used for full-length *LdCPR* cloning. The PCR was performed using a primer pair LdCPRF/LdCPRR (Table S1) which was designed based on the transcriptome sequence data (Zhu et al., 2016). The PCR products were cloned into pGEM®-T Easy Vector Systems (Promega) and sequenced. Cloning and sequence analyses of *LdCPR* gene fragments were repeated at least three times with different preparations of RNAs. Three clones from each replication were sequenced.

### 2.3. Quantitative real-time PCR (qRT-PCR)

DNase I treated total RNA was used as template for qRT-PCR, which was run in CFX96 Touch real-time PCR detection system (Bio-Rad Laboratories, Hercules, CA). Each PCR reaction (10 *μ*L final volume) contained 5 *μ*L SsoAdvanced™ Universal SYBR® Green Supermix (Bio-Rad Laboratories, Hercules, CA), 100 *ng* of cDNA, ddH2O, and 0.4 *μ*L of forward and reverse primers (Table S1, stock 10 *μ*M). The qRT-PCR condition was 95 °C for 3 min, followed by 40 cycles of 95 °C for 10 s, 55 °C for 20 s, and 72 °C for 30 s. In each plate, the positive and negative (non-template) controls and internal controls were included. A fluorescence reading determined the extension of amplification at the end of each cycle. Relative expression levels for specific genes, in relation to the most reliable reference genes *RpL4* and *Ef1α* (Zhu et al., 2016) were calculated by the 2^-△△CT^ method (Livak and Schmittgen, 2001). The PCR efficiency (95% < E < 105%) and R^2^ value (> 0.99) were considered as qualified for further analysis. Each experiment was repeated at least three times using samples from independent treatments.

### 2.4. Bioinformatics analysis

The full-length <u>o</u>pen <u>r</u>eading <u>f</u>rame (ORF) sequences of insect or spider CPR genes were extracted from GenBank (http://www.ncbi.nlm.nih.gov/) or other web sources (Table 1), and translated into protein sequences for alignment analysis using MUSCLE included in MEGA version 6.06 (http://www.megasoftware.net/) with default settings (Tamura et al., 2011). The phylogenic tree was then generated by the neighbor-joining algorithm with 2000 bootstrap replicates since the p-distance was < 0.8 in overall distance (Hall, 2011).

**Table 1.**
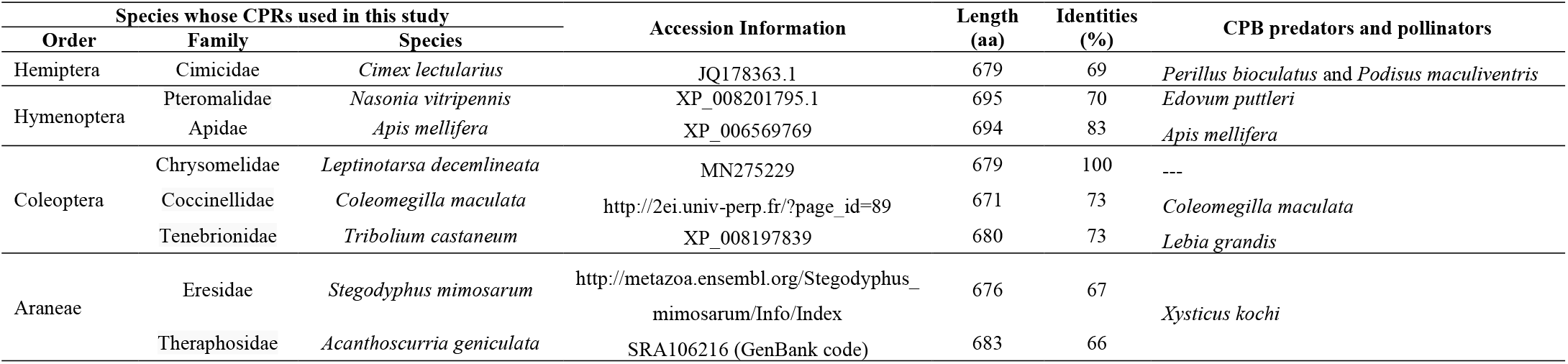
LdCPR and its homologs derived from CPB’s predators or species of predators’ closely related to CPB’s predators and pollinators.

| Species whose CPRs used in this study |  |  | Accession Information | Length<br>(aa) | Identities<br>(%) | CPB predators and pollinators |
| --- | --- | --- | --- | --- | --- | --- |
| Order | Family | Species |  |  |  |  |
| Hemiptera | Cimicidae | *Cimex lectularius* | JQ178363.1 | 679 | 69 | *Perillus bioculatus* and *Podisus maculiventris* |
| Hymenoptera | Pteromalidae | *Nasonia vitripennis* | XP_008201795.1 | 695 | 70 | *Edovum puttleri* |
|  | Apidae | *Apis mellifera* | XP_006569769 | 694 | 83 | *Apis mellifera* |
| Coleoptera | Chrysomelidae | *Leptinotarsa decemlineata* | MN275229 | 679 | 100 | --- |
|  | Coccinellidae | *Coleomegilla maculata* | http://2ei.univ-perp.fr/?page_id=89 | 671 | 73 | *Coleomegilla maculata* |
|  | Tenebrionidae | *Tribolium castaneum* | XP_008197839 | 680 | 73 | *Lebia grandis* |
| Araneae | Eresidae | *Stegodyphus mimosarum* | http://metazoa.ensembl.org/Stegodyphus_mimosarum/Info/Index | 676 | 67 | *Xysticus kochi* |
|  | Theraphosidae | *Acanthoscurria geniculata* | SRA106216 (GenBank code) | 683 | 66 |  |

### 2.5. dsRNA synthesis in bacteria

Genomic DNA (gDNA) was isolated from CPB adults using DNeasy Tissue Kit (QIAGEN) following the manufacturer’s protocol. A 244 bp fragment of the target gene was amplified by PCR with gene specific dsRNA primers (Table S1) designed based on the alignment of CPR sequences of CPB, CPB’s predators or their closely-related species, and the pollinator honey bee (Table 1). The PCR products were cloned into a T-tailed L4440 vector following the protocol described previously (Kamath and Ahringer, 2003). The single colonies of *E. coli* containing L4440 vector plus insert were inoculated into 2 x YT medium containing ampicillin and cultured overnight. The bacteria solution was diluted 100-fold with 2 x YT medium and allowed to grow to OD_600_ = 0.4. Then 1 mM (a final concentration) of IPTG was added and the culture was incubated with shaking for 5 h at 37 °C. Lastly, the solution was heat-killed by being exposed to 80 °C for 20 min and saved at −20 °C until use. To analyze the dsRNA synthesized in the bacteria HT115 (DE3), total RNA was extracted from the bacterial cells using TRIzol reagent (Invitrogen). The RNA was treated with DNase I (Invitrogen) to remove the gDNA. About 4 *μ*g of total RNA was loaded onto a 1% agarose/TBE gel and photographed.

### 2.6. dsRNA feeding bioassay

The dsRNA feeding bioassay was performed based on our previous study (Zhu et al., 2011). Briefly, one-week old female CPB adults were starved for 12 h before the initiation of the feeding assay. The 200 *μL* bacteria expressing dsRNA of *LdCPR* was spread on potato leaves in each petri dish (diameter = 8.5 cm, Fisher Scientific) to feed five female adults. The control female beetles were fed with 200 *μ*L bacteria expressing dsRNA of *GFP.* After treatment, beetles were placed in an incubator (25±1 °C, RH of 70%, L:D 16:8). The beetles were treated with dsRNA for 3 consecutive days and then allowed to feed on fresh potato leaves alone for 2 more days before being submitted to further analysis. Three biological replicates were used in both the control and treatment groups.

### 2.7. Microsome preparation and CPR enzyme activity assay

Female CPB adults treated with dsRNA of *GFP* or *LdCPR* were homogenized on ice with homogenization buffer consisting of a 50 mM Tris, 150 mM KCl, 20% Glycerol, 1 mM EDTA, 1mM PMSF, 1 mM DTT, 1 Pierce™ mini-Protease Inhibitor Tablet w/out EDTA, pH 7.6. Three beetles were pooled into one sample. Three samples were prepared for both dsRNA of *GFP* and *LdCPR* treated beetles. Each sample of homogenates were centrifuged at 12,000 g for 20 min at 4 °C. The supernatant was transferred to a SW 55Ti and centrifuged in a Beckman L8-70M ultracentrifuge at 100,000 g for 1 hr at 4 °C. Then pellets were resuspended in 50 mM Tris, 150 mM KCl, 20% Glycerol, 1 mM EDTA, 1mM PMSF, 1 mM DTT, pH 7.6, flash frozen in dry ice/ethanol bath and stored at −80 °C until later use for CPR activity assay.

CPR enzyme activity was determined by measuring the reduction of cytochrome c based on a method described by Guengerich (Guengerich et al., 2009). Briefly, microsomal protein content was determined by Bradford assay (Bradford, 1976). The enzyme assay was run in 96-well clear flat bottom plates, at 30 °C, measured in kinetic mode for 3 mins with reads at λ = 550 nm every 20 seconds on a Tecan Spark® multi-mode plate reader (Tecan U.S. Inc., Morrisville, NC 27560, USA). Reaction were performed in a volume of 200 *μ*L with a buffer consisting of 0.3 M potassium phosphate pH 7.7, 1 mM KCN, 50 *μ*M cytochrome c (from equine heart, Sigma), 4 *μ*g of protein, and 60 *μ*M NADPH (Sigma). Data in the linear range was used to calculate cytochrome c reduction activity.

### 2.8. Bioassays

Female adults were treated with serial dilutions of technical grade imidacloprid (99% active ingredient, Chem Services, West Chester, PA) prepared in acetone in the preliminary studies. A discriminating dose (causing approximately 50% mortality) of imidacloprid based our previous study was applied for the bioassays (Zhu et al., 2011; Zhu et al., 2016). Acetone only was used as control. The solution was dropped on the abdomen of the beetles (1*μ*L/drop) using a PB-600 repeating dispenser (Hamilton Co., Reno). The mortality was determined at 24 h after treatment. Mean and standard errors for each time point were obtained from at least three independent bioassays.

### 2.9. Laboratory and greenhouse assays

Four *LdCPR-* or GFP-(control) dsRNA treated virgin females and three 1-week-old males were kept in one rearing cage with new potted potato plants. Egg masses from both control and *LdCPR* dsRNA treated beetles were collected daily and the egg numbers were recorded. Then these eggs were stored in a petri dish with water-soaked filter paper and kept in an incubator (25±1 °C, RH of 70%, L:D=16:8). The hatching percentage of eggs from dsRNA of *LdCPR* treated and control females were calculated. Four replicates per treatment were performed.

### 2.10. Embryo fixing and DAPI staining

*L. decemlineata* eggs were collected in 5 days after laid. The eggs were dechlorinated in 25% bleach for 2 min and rinsed well in deionized water. Then the CPB embryos were carefully transferred into a glass scintillation vial containing fixing solution and had been fixed for 20 min on a shaking platform as previous study described (Bitra and Palli, 2010). The embryos were devitellinized by adding 8 ml of methanol and then used for nuclear staining. Nuclear staining was done with DAPI (4’,6-diamidino-2-phenylindole, dihydrochloride, Sigma) at 1:1000 dilution of 1 mg/ml stock for 10 min. The stained embryos were mounted on a slide for photography using Olympus 1×71 Inverted Research Microscope.

### 2.11. Statistical analysis

Statistical significance of the gene expression among samples was calculated using one-way analysis of variation (ANOVA) followed by Duncan multiple mean separation techniques with SAS v9.2 software, and the statistical significance between two samples was calculated using a Student’s t-test (two-sample comparison). Different letters (i.e., a, b and c) were used to indicate significant difference (*P* < 0.05) among relative expression within samples.

## 3. Results

### 3.1. LdCPR cloning and developmental expression pattern

Based on our previous transcriptome and annotation data (Zhu et al., 2016), only one CPR candidate gene was identified. With the available gene information, specific primers were designed to amplify and the full-length *LdCPR* gene was cloned from the cDNA sample of the imidacloprid resistant *L. decemlineata* strain. The 2040 bp cDNA sequence was submitted to the GenBank database (accession number MN275229).

The mRNA levels of *LdCPR* in the one day-old eggs (E1), the five day-old eggs (E5), 1^st^ to 4^th^ instar larvae (L1 – L4), pupae (P), and one week-old adult female (AF) and adult male (AM) of imidacloprid resistant *L. decemlineata* were quantified. The highest expression level was detected in E1. The second highest expression level was observed in L2. The lowest expression level of *LdCPR* was detected in the P stage (Fig. 1).

**Fig. 1.**
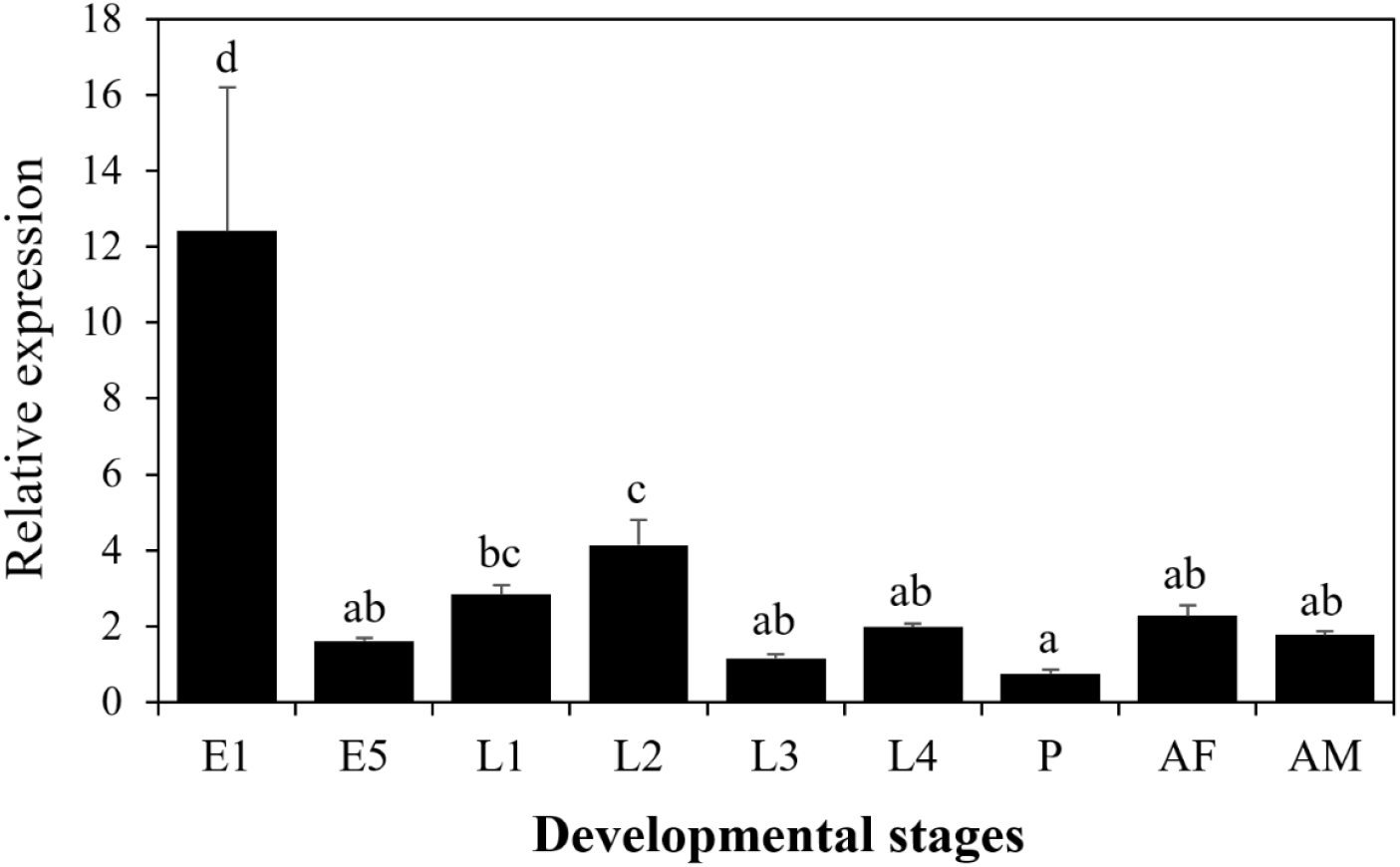
Developmental expression pattern of *LdCPR* determined by qRT-PCR in the imidacloprid resistant *L. decemlineata.* Seven developmental insect stages were used, including one day-old eggs (E1), five day-old eggs (E5), 1^st^ through 4^th^ instar larvae (L1, L2, L3, and L4, respectively), pupae (P), 1-week-old adult females (AF), and 1-week-old adult males (AM). The relative mRNA levels were shown as a ratio in comparison with the levels of *RpL4* and *Ef1α* mRNA. The data shown are mean + SEM (n = 6). Statistical significance of the gene expression among stages was calculated using ANOVA followed by Duncan multiple mean separation techniques. There was no significant difference (*P* < 0.05) among expressions with the same alphabetic letter (i.e. a, b and c).

### 3.2. LdCPR dsRNA synthesis strategy

A comprehensive dsRNA synthesis strategy was used to minimize the effect of *LdCPR* dsRNA on nontarget organisms including predators or the closely related species of predators of *L. decemlineata* and surrogate species of pollinators such as honeybee. We chose the most closely related species with available sequence information: *Cimex lectularius* (same family as *L. decemlineata’*s predators *Perillus bioculatus and Podisus maculiventris), Nasonia vitripennis* (same family as *L. decemlineata’*s predator *Edovum puttleri), Apis mellifera, Coleomegilla maculate, Tribolium castaneum* (same family as *L. decemlineata’*s predator *Lebia grandis), Stegodyphus mimosarum* and *Acanthoscurria geniculata* (same order as *L. decemlineata*’s predator *Xysticus kochi*) (Table 1). The sequences of CPR from these species were obtained from GenBank or other web resources (Table 1). A phylogenetic tree for these CPR sequences was generated by MEGA 6.06 using the neighbor-joining algorithm with amino acid sequences of all 8 species (Fig. 2A). Results showed that the CPRs from the same order were clustered into the same branch. LdCPR originated from the same evolutionary root as TcCPR with a bootstrap value of 65 (Fig. 2A).

**Fig. 2.**
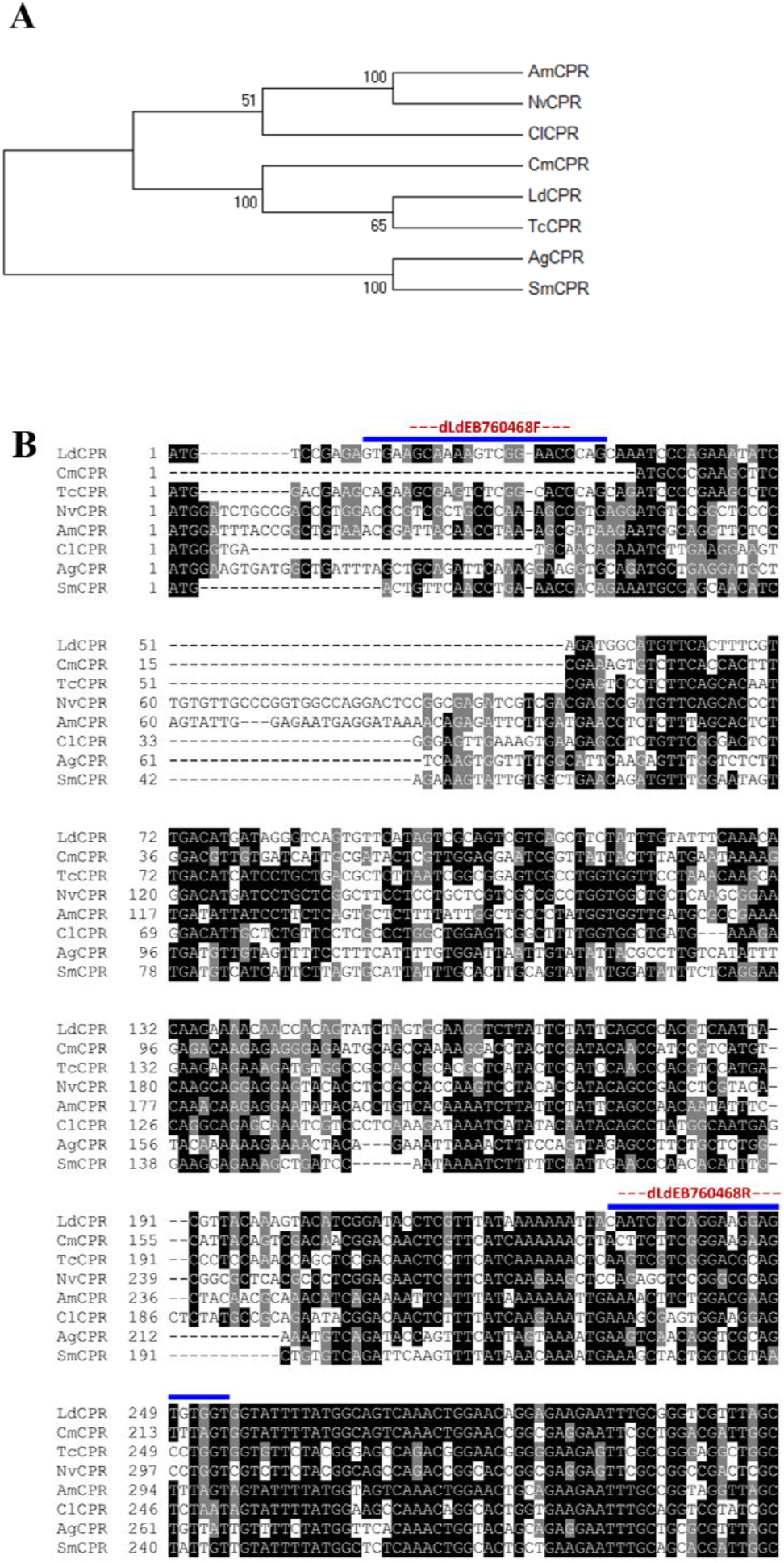
*LdCPR* dsRNA synthesis strategy. A, Phylogenetic relationship of LdCPR with its orthologs in the predators or closely related species to predators of *L. decemlineata* and a pollinator. B, sequence alignment of *LdCPR* in selected species. Darker colors indicate higher degree of sequence similarity shared among species selected. The blue lines highlight the designed primer sequence regions.

The DNA sequence corresponding to *LdCPR* dsRNA exhibited a low level of sequence similarity to the CPR genes of nontarget species (Fig. 2B). Specifically, within the DNA region that was selected to synthesize *LdCPR* dsRNA, there was not a sequence fragment over 19-nt long (Whyard et al., 2009; Bachman et al., 2013) identical to the sequences of any other species compared (Fig. 2B), including the most closely related species, *T. castaneum.* Thus, the likelihood of that the dsRNA we developed to target *LdCPR* gene have nontarget effect on the genes of other species was reduced. As shown in the Fig. S1, the dsRNA of *LdCPR* was successfully synthesized in the HT115 *E. coli* cell line.

### 3.3. LdCPR RNAi decreased imidacloprid resistance

The knockdown efficiency of *LdCPR* RNAi on female imidacloprid resistant beetles was measured by detecting changes in mRNA levels and enzymatic activities on the 6^th^ day after the initiation of feeding RNAi, using qRT-PCR and CPR enzyme activity assay, respectively. The mRNA level of *LdCPR* decreased by 82% on the 6^th^ day after feeding with dsRNA of *LdCPR* compared with the control (Fig. 3A). CPR enzyme activity toward cytochrome c reduction decreased by about 50% on the 6^th^ day after feeding with dsRNA of *LdCPR* compared with the control (Fig. 3B). These results suggest LdCPR was successfully knocked down in both mRNA and protein levels. The following bioassay with imidacloprid showed that knockdown of *LdCPR* in female beetles significantly increased the susceptibility to imidacloprid (Fig. 3C). The percent survivals of dsRNA treated imidacloprid resistant female beetles at 1 *μ*g imidacloprid on the 6^th^ day after dsRNA injection were 63% and 10% in the control and *LdCPR-KD* beetles, respectively (Fig. 3C).

**Fig. 3.**
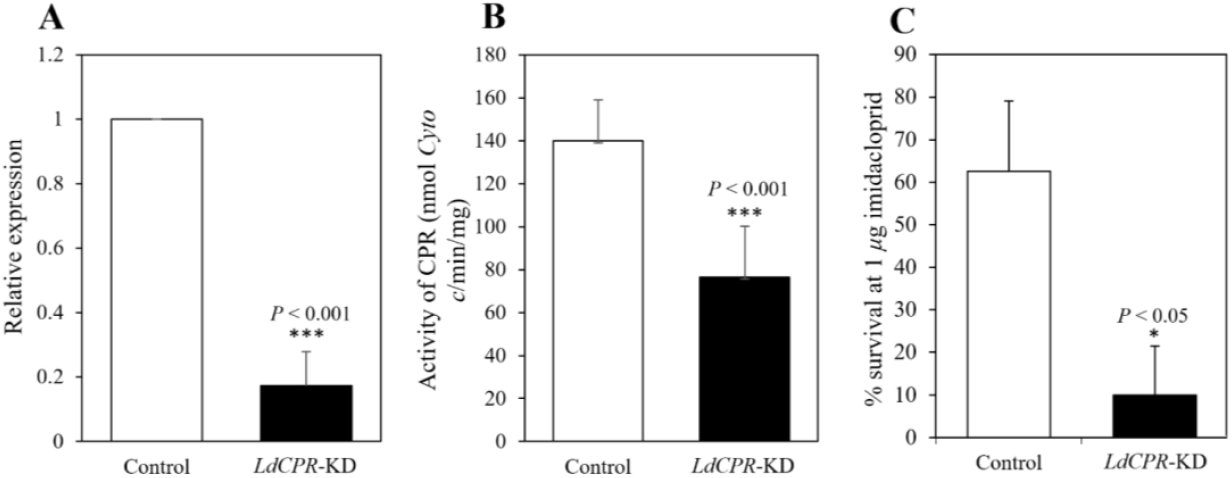
Knockdown (KD) of LdCPR reduced the resistance of *L. decemlineata* to imidacloprid. Beetles fed with *LdCPR* dsRNA (LdCPR-KD) and fed with *GFP* dsRNA (Control) were compared. A, Expression of *LdCPR* at the mRNA level was evaluated by qRT-PCR. B, The enzymatic activities of LdCPR after dsRNA ingestion were analyzed by CPR enzyme assay. C, Insect survival rate on the 6^th^ day after dsRNA injection. The mortality was recorded after 24 h exposure to imidacloprid (3 replicates, 30 individuals for each replicate). The mortality in acetone controls was zero. The difference between control (*GFP*) and *LdCPR*-KD imidacloprid resistant beetles was analyzed by Student’s *t*-test.

### 3.4. Parental RNAi efficiency in eggs

The mRNA levels of *LdCPR* in the eggs laid by female beetles (Fig. 4A) fed with dsRNA of *GFP* and *LdCPR* were detected by qRT-PCR. For 3-day old eggs, the *LdCPR* expression level decreased by 66% in eggs laid by beetles fed with dsRNA of *LdCPR* compared with eggs laid by beetles fed with dsRNA of *GFP* (Fig. 4B), suggesting the RNAi effects were successfully transferred to the next generation.

**Fig. 4.**
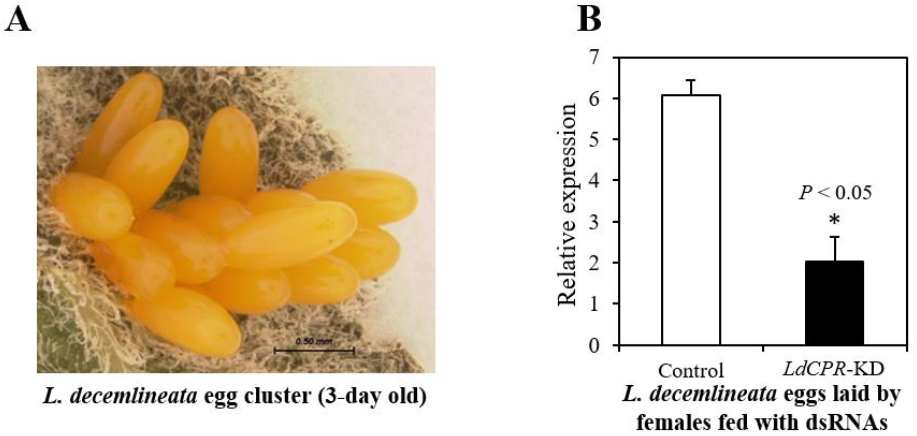
Silencing *LdCPR* in susceptible *L. decemlineata* female adults causes parental RNAi in eggs. A, An egg cluster laid by a female *L. decemlineata.* B, Relative mRNA levels of *LdCPR* in eggs laid by *GFP* (control) or *LdCPR* (*LdCPR*-KD) dsRNA-treated female adults. Three replicates were performed, each including 30 eggs of 3 different egg clusters laid by one female. The difference between control (*GFP*) and *LdCPR*-KD imidacloprid resistant beetles was analyzed by Student’s *t*-test.

### 3.5. Feeding RNAi effects on female fecundity and egg hatching

As shown in the Fig. 5, knockdown of *LdCPR* led to significantly decreased egg number laid by each female on each day. On average, each female fed with dsRNA of GFP laid about 21 eggs each day. After knockdown of *LdCPR*, each female only laid about 15 eggs each day (Fig. 5C). In addition, silencing of *LdCPR* caused enhance egg mortality (Fig. 5D). In the control beetles, the percentage of egg hatching was about 73%. However, after knockdown of the *LdCPR* from the beetles, the percentage of egg hatching decreased down to just 38% (Fig. 5D).

**Fig. 5.**
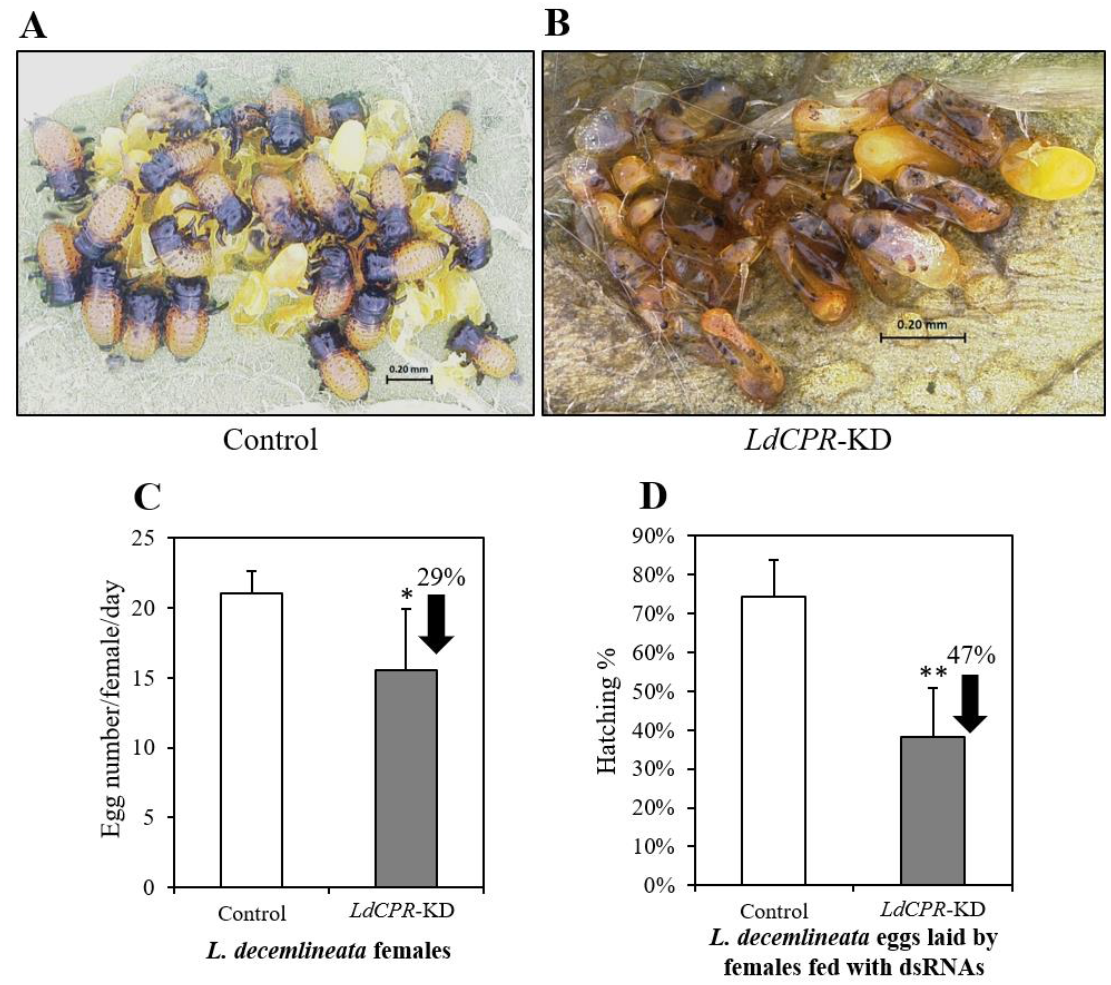
RNAi of *LdCPR* in *L. decemlineata* beetles lead to reduced female fecundity and enhanced egg mortality. A, Eggs hatching in the control (laid by females fed with dsRNA of *GFP).* B, Eggs failed in hatching in *LdCPR*-KD females. C, Average egg number laid by each female fed with dsRNA of *GFP* or *LdCPR* in each day. D, Percentage of hatching eggs laid by females fed with dsRNA of *GFP* and *LdCPR.* The differences between control (*GFP*) and *LdCPR*-KD beetles were analyzed by Student’s *t*-test. * *P* < 0.05; ** *P* < 0.01.

### 3.6. Parental RNAi effects on embryo development

At the temperature of 25±1 °C, it took about 5 days for an egg to finish development and hatch to the 1^st^ instar larva. As shown in the Fig. 6A, the eggs laid by control beetles (treated with dsRNA of *GFP*) developed to the fully extended germband stage and was ready to hatch. After knockdown of the *LdCPR*, the embryo development was stuck at the early germband stage (Fig. 6B) and could not complete the normal development, eventually caused embryo death (Fig. 5B).

**Fig. 6.**
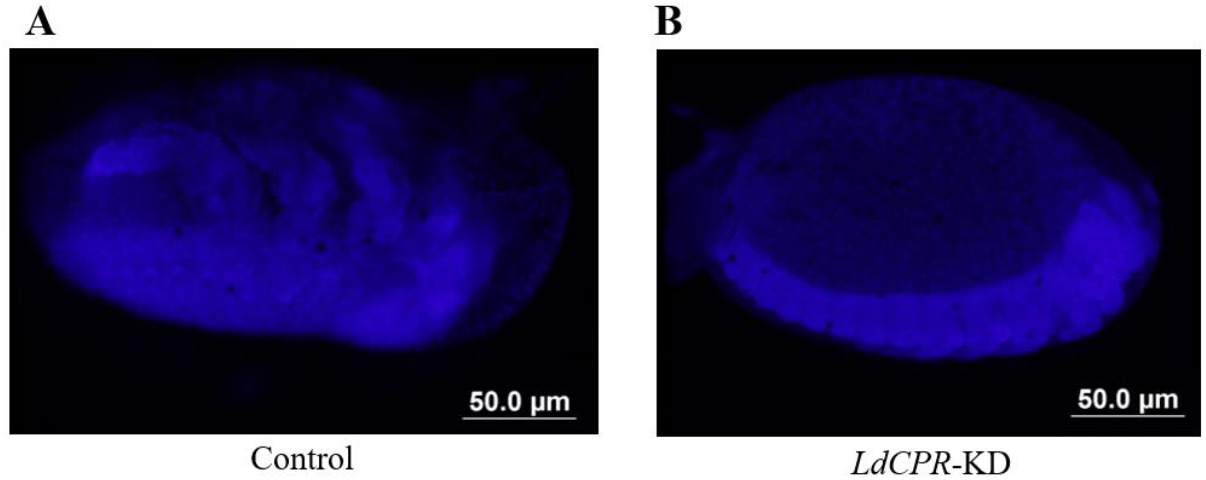
RNAi of *LdCPR* in *L. decemlineata* adults caused their egg embryos to be stuck in the 5^th^ day of development. A, Embryo produced by beetle fed on *GFP* dsRNA (control). B, Embryo produced by beetle fed on *LdCPR*-KD dsRNA. Embryos were stained with DAPI and photographed.

## 4. Discussions

During the past two decades, RNAi has been explored as a new and complementary pest control tactic to meet increasing demand for sustainable and environmentally benign pest control strategies (Swevers et al., 2013; Zhu et al., 2014; Zotti et al., 2018; Ma et al., 2019). Compared with traditional chemical control methods, RNAi-based biopesticides have the advantage in their unique mode of action being dependent on the target gene sequences. However, unintentional off-target effects in target organisms and silencing effects in non-target organisms occur more often than expected (Qiu et al., 2005; Baum et al., 2007; Lundgren and Duan, 2013; Zhu et al., 2014). Careful experiment design with bioinformatics analysis is one feasible approach to minimize nontarget species-specific effects with RNAi (Bachman et al., 2013). Whyard et al. revealed that once dsRNA target regions of genes have no shared 19-21 nt sequence with other nontarget species tested, even closely related species of the same genus can be selectively controlled (Whyard et al., 2009). In our current study we designed dsRNA primers based on this principle, by sequence alignment of *LdCPR* with other *CPR* sequences from seven species (Table 1; Fig. 2B). From the genome/transcriptome-sequenced insect species, we selected the ones that are phylogenetically closely related to CPB’s predators (i.e., belonging to same family), and *A. mellifera* which is a well-known beneficial insect species (Romeis et al., 2008; Velez et al., 2016b), for analysis. The purpose of this dsRNA synthesis strategy was to reduce the sequence similarity with nontarget species and offer a high degree of species specificity for *LdCPR* RNAi.

Imidacloprid is the most extensively used neonicotinoid insecticide (IRAC group 4A), which plays a vital role in preventing crops from many major agricultural pests, such as sucking insect pests, coleopteran pests, and some micro lepidoptera (Elbert et al., 2008). Since firstly launched in 1991, resistance to imidacloprid has been reported in numerous insect species including western flower thrips (Zhao et al., 1995), tobacco whiteflies (Elbert and Nauen, 2000), Colorado potato beetle (Zhao et al., 2000), fruit fly (Daborn et al., 2001), brown planthopper (Zewen et al., 2003), aphids (Nauen and Denholm, 2005), and small brown planthopper (Sanada-Morimura et al., 2011). The mechanisms of imidacloprid resistance in insects have been linked with a mutation in the target protein nicotinic acetylcholine receptor (nAChR) (Liu et al., 2005) and more commonly cytochrome P450-mediated detoxification (Karunker et al., 2008; Puinean et al., 2010; Hoi et al., 2014; Elzaki et al., 2017; Sun et al., 2018; Chen et al., 2019). In *L. decemlineata*, biochemical, physiological, and molecular biological studies showed that cytochrome-P450 mediated detoxification is the most common mechanism in neonicotinoid resistance (Nauen and Denholm, 2005; Alyokhin et al., 2008; Zhu et al., 2016), although the contributions from other mechanisms cannot be completely ruled out (Clements et al., 2016; Clements et al., 2017). A recent RNA-seq study identified several cytochrome P450 genes overexpressed in imidacloprid resistant populations (Clements et al., 2016). Then the following functional genomics study revealed that one cytochrome P450 (Comp 115309) may play an important role in imidacloprid resistance (Clements et al., 2017). In our current study, after knocking down the cytochrome P450 partner, *LdCPR*, cytochrome P450s’ function interrupted, which caused the imidacloprid resistance decreased significantly (Fig. 3), indicating cytochrome P450-mediated detoxification is responsible for imidacloprid resistance in CPB.

Inactivating microsomal P450s (Henderson et al., 2003) through the inhibition or silencing of CPR, offers a unique way to decipher contributions of P450s in adaptation to one or multiple pesticides. Recent studies showed that silencing CPR in several insect species increased susceptibilities to pyrethroids, neonicotinoids, organophosphates, carbamates, or avermectins, suggesting the profound roles of cytochrome P450s in insecticide adaptation (Lycett et al., 2006; Zhu et al., 2012; Huang et al., 2015; Liu et al., 2015; Wang et al., 2016; Zhang et al., 2016; Zhang et al., 2017; Jing et al., 2018). Most recently, a study in a polyphagous agricultural pest, *Tetranychus urticae* found that downregulation of CPR led to increased susceptibilities to multiple acaricides commonly used for pest control, including abamectin, bifenthrin and fenpyroximate with distinct modes of action (Adesanya et al., 2020). Here we showed that studies on CPR inhibition not only facilitate our understanding on the contribution of cytochrome P450-mediated detoxification to pesticide adaptation but also provide a unique path for practically delaying the development of pesticide resistance in the field. Due to the essential role of CPR in electron transfer to numerous microsomal electron acceptors, even a single nonsynonymous mutation has potential to significantly alter activities of CPR and certain cytochrome P450s. In human, approximately 2000 single nucleotide polymorphisms (SNPs) including more than 150 nonsynonymous mutations have been identified on human POR gene (equivalent to CPR). Many of these mutations are tightly associated with human diseases, such as Antley-Bixler Syndrome and disordered steroidogenesis (Waskell and Kim, 2015). In the mosquito *Anopheles minimus*, nonsynonymous mutations L86F, L219F, C427R were found to increase FMN binding, enzyme stability, and cytochrome c reduction and potentially enhance the activity of a pyrethroid resistance-associate P450 (Sarapusit et al., 2008; Sarapusit et al., 2013). In the bird cherry-oat aphid *Rhopalosiphum padi*, 334 SNPs including 194 nonsynonymous mutations were detected from 11 geographic populations which may be associated with P450-mediated insecticide resistance in *R. padi.* One mutation P484S located in the FAD-binding region of CPR was found in more than 35% of individual aphids across populations (Wang et al., 2016). Most recently, two bulked segregant analysis genomic mapping studies in acaricide resistant *T. urticae* strains identified *TuCPR* as a potential genomic locus that was associated with resistance to spirodiclofen, pyridaben and tebufenpyrad. A nonsynonymous mutation, D384Y, in TuCPR was detected in acaricide resistant *T. urticae* strains (Snoeck et al., 2019; Wybouw et al., 2019).

Besides metabolism of xenobiotics, cytochrome P450s are also involved in endogenous compounds biosynthesis pathways including juvenile hormone and molting hormone (Helvig et al., 2004a; Rewitz et al., 2007). For example, CYP15A1 catalyzes epoxidation of methyl farnesoate to juvenile hormone III in cockroach corpora allata (Helvig et al., 2004a). Insect genomes contain single orthologs of Halloween genes that are a group of P450 enzymes required for the biosynthesis pathway of molting hormone, 20-hydroxyecdysone (20E) (Rewitz et al., 2007). These Halloween genes include microsomal P450s *Phantom (CYP306A1)* and *spook/spookier (CYP307A1/A2)* as well as several mitochondrial P450s (Rewitz et al., 2007). These P450s are expressed in embryos, larval stages, and adult ovaries, suggesting their important functions in embryonic development, metamorphosis, and reproduction. Suppression of the *LdCPR* expression led to significant decrease of fecundity of female beetles (29% less eggs/day) (Fig. 5C). It is possible that silencing *LdCPR* disrupted the functions of the P450s involved in biosynthesis of hormones that are critical for insect reproduction. After knocking down *LdCPR* in female adults, the hatching rate of their eggs decreased dramatically (Fig. 5D). This suggested that *LdCPR* RNAi may influence the biosynthesis of 20E in female ovaries and decrease the amount of 20E molecules that are vertically transmitted to the offspring’s embryos (Truman and Riddiford, 1999). Alternatively, the 20E biosynthesis in the embryos could be suppressed by *LdCPR* dsRNAs that are maternally deposited by female adults. In addition, it was shown that inhibition of CPR in silkworm embryos caused a reduction of microsomal ecdysone 20-hydroxylase activity (Horike et al., 2000). In mouse, an abolished expression of the CPR gene in the early embryonic stage caused multiple developmental defects and some embryonic lethality (Shen et al., 2002). In human, mutations in CPR led to disordered biosynthesis of steroid hormones, which is required for development, reproduction, and stress responses (Fluck et al., 2004).

In our study, the bacteria-expressed dsRNA-mediated feeding RNAi of *LdCPR* in female adults caused systemic knock down in the expression of the gene coding for *LdCPR* in both adults and their eggs (Figs. 3A and 4B). This is the first report of parental RNAi effect in the CPB, although such a phenomenon has been observed in *Tribolium castaneum* (Bucher et al., 2002), *Diabrotica virgifera virgifera* (Velez et al., 2016a), and *Euschistus heros* (Fishilevich et al., 2016). Knowledge of parental RNAi response will benefit the management of CPB with RNAi-based biopesticides for crop protection.

## Acknowledgements

This project was supported by a faculty start-up fund from Pennsylvania State University, and the USDA National Institute of Food and Federal Appropriations under Hatch Project #PEN04609 and Accession #1010058 (to F.Z.). M.W. was supported by the Education Department of Fujian Province of China. We are grateful Drs. Douglas B. Walsh and Laura C. Lavine for their support and Dr. James H. Tumlinson for sharing lab equipment with us.

## Supplementary data

**Table 1S.**
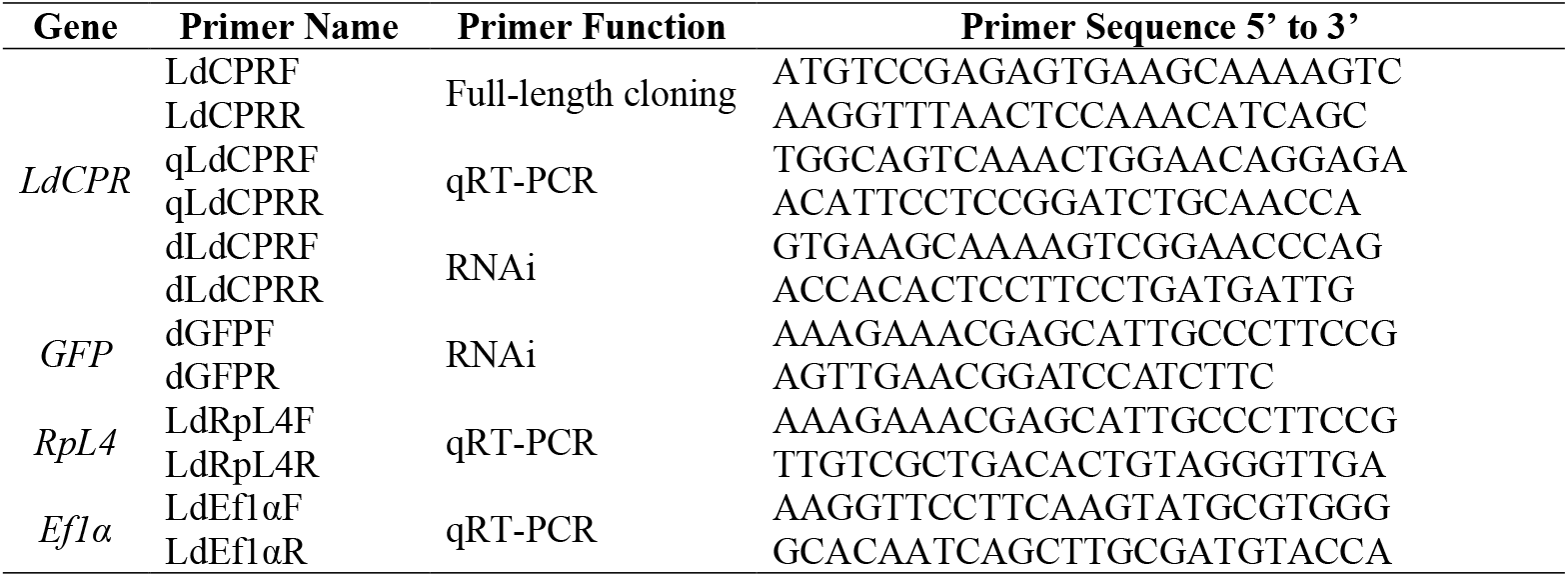
Primers used for gene cloning, qRT-PCR and dsRNA synthesis.

**Fig. 1S.**
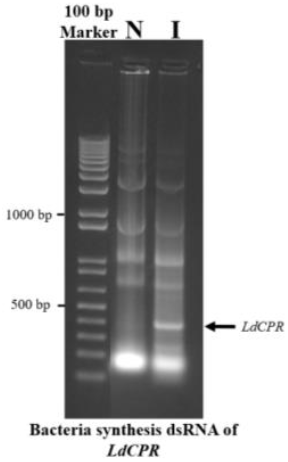
Identification of dsRNA of *LdCPR* produced in bacteria. Total RNA from the non-induced bacteria (N) and IPTG-induced bacteria (I) was extracted and run on agarose gel. The arrow pointed to the band of *LdCPR* dsRNA produced in IPTG-induced bacteria.

## Notes

### Competing Interest Statement

The authors have declared no competing interest.

## References

Adesanya, A.W., Cardenas, A., Lavine, M.D., Walsh, D.B., Lavine, L.C., Zhu, F., 2020. RNA interference of NADPH-Cytochrome P450 reductase increases susceptibilities to multiple acaricides in *Tetranychus urticae*. Pesticide Biochemistry and Physiology 165, 104550.

Alyokhin, A., Baker, M., Mota-Sanchez, D., Dively, G., Grafius, E., 2008. Colorado potato beetle resistance to insecticides. American Journal of Potato Research 85, 395–413.

Alyokhin, A., Mota-Sanchez, D., Baker, M., Snyder, W.E., Menasha, S., Whalon, M., Dively, G., Moarsi, W.F., 2015. The Red Queen in a potato field: integrated pest management versus chemical dependency in Colorado potato beetle control. Pest Management Science 71, 343–356.

Bachman, P.M., Bolognesi, R., Moar, W.J., Mueller, G.M., Paradise, M.S., Ramaseshadri, P., Tan, J., Uffman, J.P., Warren, J., Wiggins, B.E., Levine, S.L., 2013. Characterization of the spectrum of insecticidal activity of a double-stranded RNA with targeted activity against Western Corn Rootworm (*Diabrotica virgifera virgifera* LeConte). Transgenic Research 22, 1207–1222.

Balabanidou, V., Kampouraki, A., MacLean, M., Blomquist, G.J., Tittiger, C., Juarez, M.P., Mijailovsky, S.J., Chalepakis, G., Anthousi, A., Lynd, A., Antoine, S., Hemingway, J., Ranson, H., Lycett, G.J., Vontas, J., 2016. Cytochrome P450 associated with insecticide resistance catalyzes cuticular hydrocarbon production in *Anopheles gambiae*. Proceedings of the National Academy of Sciences of the United States of America 113, 9268–9273.

Baum, J.A., Bogaert, T., Clinton, W., Heck, G.R., Feldmann, P., Ilagan, O., Johnson, S., Plaetinck, G., Munyikwa, T., Pleau, M., Vaughn, T., Roberts, J., 2007. Control of coleopteran insect pests through RNA interference. Nature Biotechnology 25, 1322–1326.

Bitra, K., Palli, S.R., 2010. The members of bHLH transcription factor superfamily are required for female reproduction in the red flour beetle, *Tribolium castaneum*. Journal of Insect Physiology 56, 1481–1489.

Bradford, M.M., 1976. A rapid and sensitive method for the quantitation of microgram quantities of protein utilizing the principle of protein-dye binding. Analytical Biochemistry 72, 248–254.

Bucher, G., Scholten, J., Klingler, M., 2002. Parental RNAi in *Tribolium* (Coleoptera). Current Biology 12, R85–86.

Cariño, F., Koener, J.F., Plapp Jr., F.W., Feyereisen, R., 1992. Expression of the cytochrome P450 gene *CYP6A1* in the housefly, *Musca domestica*. in: Mullin, C.A., Scott, J.G. (Eds.). Molecular Mechanisms of Insecticide Resistance, pp. 31–40.

Chen, C., Shan, T., Liu, Y., Wang, C., Shi, X., Gao, X., 2019. Identification and functional analysis of a cytochrome P450 gene involved in imidacloprid resistance in *Bradysia odoriphaga* Yang et Zhang. Pesticide Biochemistry and Physiology 153, 129–135.

Clements, J., Schoville, S., Peterson, N., Huseth, A.S., Lan, Q., Groves, R.L., 2017. RNA interference of three up-regulated transcripts associated with insecticide resistance in an imidacloprid resistant population of *Leptinotarsa decemlineata*. Pesticide Biochemistry and Physiology 135, 35–40.

Clements, J., Schoville, S., Peterson, N., Lan, Q., Groves, R.L., 2016. Characterizing molecular mechanisms of imidacloprid resistance in select populations of *Leptinotarsa decemlineata* in the central sands region of Wisconsin. PloS One 11, e0147844.

Coon, M.J., Vaz, A.D., Bestervelt, L.L., 1996. Cytochrome P450 2: peroxidative reactions of diversozymes. FASEB Journal 10, 428–434.

Daborn, P., Boundy, S., Yen, J., Pittendrigh, B., ffrench-Constant, R., 2001. DDT resistance in *Drosophila* correlates with Cyp6g1 over-expression and confers cross-resistance to the neonicotinoid imidacloprid. Molecular Genetics and Genomics: MGG 266, 556–563.

David, J.P., Ismail, H.M., Chandor-Proust, A., Paine, M.J., 2013. Role of cytochrome P450s in insecticide resistance: impact on the control of mosquito-borne diseases and use of insecticides on Earth. Philosophical Transactions of the Royal Society of London B Biological Sciences 368, 20120429.

Elbert, A., Haas, M., Springer, B., Thielert, W., Nauen, R., 2008. Applied aspects of neonicotinoid uses in crop protection. Pest Management Science 64, 1099–1105.

Elbert, A., Nauen, R., 2000. Resistance ofBemisia tabaci (Homoptera:Aleyrodidae) to insecticides in southern Spain with special reference to neonicotinoids. Pest Management Science 56, 60–64.

Elzaki, M.E.A., Miah, M.A., Wu, M., Zhang, H., Pu, J., Jiang, L., Han, Z., 2017. Imidacloprid is degraded by CYP353D1v2, a cytochrome P450 overexpressed in a resistant strain of *Laodelphax striatellus*. Pest Management Science 73, 1358–1363.

Feyereisen, R., 1999. Insect P450 enzymes. Annual review of entomology 44, 507–533.

Feyereisen, R., 2005. Insect cytochrome P450. in: Gilbert, L.I., Latrou, K., Gill, S.S. (Eds.). Comprehensive Molecular Insect Science. Elsevier, Oxford, UK, pp. 1–77.

Feyereisen, R., 2012. Insect CYP genes and P450 enzymes. in: Gilbert, L.I. (Ed.). Insect Molecular Biology and Biochemistry. Elsevier, pp. 236–316.

Feyereisen, R., Koener, J.F., Farnsworth, D.E., Nebert, D.W., 1989. Isolation and sequence of cDNA encoding a cytochrome P-450 from an insecticide-resistant strain of the house fly, *Musca domestica*. Proceedings of the National Academy of Sciences of the United States of America 86, 1465–1469.

Fishilevich, E., Velez, A.M., Khajuria, C., Frey, M.L., Hamm, R.L., Wang, H., Schulenberg, G.A., Bowling, A.J., Pence, H.E., Gandra, P., Arora, K., Storer, N.P., Narva, K.E., Siegfried, B.D., 2016. Use of chromatin remodeling ATPases as RNAi targets for parental control of western corn rootworm (*Diabrotica virgifera virgifera*) and Neotropical brown stink bug *(Euschistus heros)*. Insect Biochemistry and Molecular Biology 71, 58–71.

Fluck, C.E., Tajima, T., Pandey, A.V., Arlt, W., Okuhara, K., Verge, C.F., Jabs, E.W., Mendonca, B.B., Fujieda, K., Miller, W.L., 2004. Mutant P450 oxidoreductase causes disordered steroidogenesis with and without Antley-Bixler syndrome. Nature Genetics 36, 228–230.

Guengerich, F.P., Martin, M.V., Sohl, C.D., Cheng, Q., 2009. Measurement of cytochrome P450 and NADPH-cytochrome P450 reductase. Nature Protocols 4, 1245–1251.

Hall, B.G., 2011. Phylogenetic trees made easy: a how-to manual Sinauer Associates.

Helvig, C., Koener, J.F., Unnithan, G.C., Feyereisen, R., 2004a. CYP15A1, the cytochrome P450 that catalyzes epoxidation of methyl farnesoate to juvenile hormone III in cockroach corpora allata. Proceedings of the National Academy of Sciences of the United States of America 101, 4024–4029.

Helvig, C., Tijet, N., Feyereisen, R., Walker, F.A., Restifo, L.L., 2004b. Drosophila melanogaster CYP6A8, an insect P450 that catalyzes lauric acid (omega-1)-hydroxylation. Biochemical and Biophysical Research Communications 325, 1495–1502.

Henderson, C.J., Otto, D.M., Carrie, D., Magnuson, M.A., McLaren, A.W., Rosewell, I., Wolf, C.R., 2003. Inactivation of the hepatic cytochrome P450 system by conditional deletion of hepatic cytochrome P450 reductase. The Journal of Biological Chemistry 278, 13480–13486.

Hoi, K.K., Daborn, P.J., Battlay, P., Robin, C., Batterham, P., O’Hair, R.A., Donald, W.A., 2014. Dissecting the insect metabolic machinery using twin ion mass spectrometry: a single P450 enzyme metabolizing the insecticide imidacloprid *in vivo*. Analytical Chemistry 86, 3525–3532.

Horike, N., Takemori, H., Nonaka, Y., Sonobe, H., Okamoto, M., 2000. Molecular cloning of NADPH-cytochrome P450 oxidoreductase from silkworm eggs. Its involvement in 20-hydroxyecdysone biosynthesis during embryonic development. European Journal of Biochemistry 267, 6914–6920.

Huang, Y., Lu, X.P., Wang, L.L., Wei, D., Feng, Z.J., Zhang, Q., Xiao, L.F., Dou, W., Wang, J.J., 2015. Functional characterization of NADPH-cytochrome P450 reductase from *Bactrocera dorsalis*: possible involvement in susceptibility to malathion. Scientific Reports 5, 18394.

Jing, T.X., Tan, Y., Ding, B.Y., Dou, W., Wei, D.D., Wang, J.J., 2018. NADPH-Cytochrome P450 Reductase mediates the resistance of *Aphis (Toxoptera) citricidus* (Kirkaldy) to Abamectin. Frontiers in Physiology 9, 986.

Kadoić Balaško, M., Mikac, K.M., Bažok, R., Lemic, D., 2020. Modern techniques in Colorado potato beetle (*Leptinotarsa decemlineata* Say) control and resistance management: history review and future perspectives. Insects 11.

Kamath, R.S., Ahringer, J., 2003. Genome-wide RNAi screening in *Caenorhabditis elegans*. Methods (San Diego, Calif.) 30, 313–321.

Karunker, I., Benting, J., Lueke, B., Ponge, T., Nauen, R., Roditakis, E., Vontas, J., Gorman, K., Denholm, I., Morin, S., 2008. Over-expression of cytochrome P450 CYP6CM1 is associated with high resistance to imidacloprid in the B and Q biotypes of *Bemisia tabaci* (Hemiptera: Aleyrodidae). Insect Biochemistry and Molecular Biology 38, 634–644.

Krieger, R.I., Feeny, P.P., Wilkinson, C.F., 1971. Detoxication enzymes in the guts of caterpillars: an evolutionary answer to plant defenses? Science (New York, N.Y.) 172, 579–581.

Liu, S., Liang, Q.M., Zhou, W.W., Jiang, Y.D., Zhu, Q.Z., Yu, H., Zhang, C.X., Gurr, G.M., Zhu, Z.R., 2015. RNA interference of NADPH-cytochrome P450 reductase of the rice brown planthopper, *Nilaparvata lugens*, increases susceptibility to insecticides. Pest Management Science 71, 32–39.

Liu, Z., Williamson, M.S., Lansdell, S.J., Denholm, I., Han, Z., Millar, N.S., 2005. A nicotinic acetylcholine receptor mutation conferring target-site resistance to imidacloprid in *Nilaparvata lugens* (brown planthopper). Proceedings of the National Academy of Sciences of the United States of America 102, 8420–8425.

Livak, K.J., Schmittgen, T.D., 2001. Analysis of relative gene expression data using real-time quantitative PCR and the 2(T)(-Delta Delta C) method. Methods 25, 402–408.

Lundgren, J.G., Duan, J.J., 2013. RNAi-based insecticidal crops: potential effects on nontarget species. BioScience 63, 657–665.

Lycett, G.J., McLaughlin, L.A., Ranson, H., Hemingway, J., Kafatos, F.C., Loukeris, T.G., Paine, M.J., 2006. Anopheles gambiae P450 reductase is highly expressed in oenocytes and in vivo knockdown increases permethrin susceptibility. Insect Molecular Biology 15, 321–327.

Ma, M., He, W., Xu, S., Xu, L., Zhang, J., 2019. RNA interference in Colorado potato beetle (Leptinotarsa decemlineata): a potential strategy for pest control. Journal of Integrative Agriculture 18, 2–11.

Maibeche-Coisne, M., Nikonov, A.A., Ishida, Y., Jacquin-Joly, E., Leal, W.S., 2004. Pheromone anosmia in a scarab beetle induced by in vivo inhibition of a pheromone-degrading enzyme. Proceedings of the National Academy of Sciences of the United States of America 101, 11459–11464.

Nauen, R., Denholm, I., 2005. Resistance of insect pests to neonicotinoid insecticides: current status and future prospects. Archives of Insect Biochemistry and Physiology 58, 200–215.

Palli, S.R., 2014. RNA interference in Colorado potato beetle: steps toward development of dsRNA as a commercial insecticide. Current Opinion in Insect Science 6, 1–8.

Porter, T.D., Kasper, C.B., 1986. NADPH-cytochrome P-450 oxidoreductase: flavin mononucleotide and flavin adenine dinucleotide domains evolved from different flavoproteins. Biochemistry 25, 1682–1687.

Puinean, A.M., Foster, S.P., Oliphant, L., Denholm, I., Field, L.M., Millar, N.S., Williamson, M.S., Bass, C., 2010. Amplification of a cytochrome P450 gene is associated with resistance to neonicotinoid insecticides in the aphid *Myzus persicae*. PLoS Genetics 6, e1000999.

Qiu, S., Adema, C.M., Lane, T., 2005. A computational study of off-target effects of RNA interference. Nucleic Acids Research 33, 1834–1847.

Qiu, Y., Tittiger, C., Wicker-Thomas, C., Le Goff, G., Young, S., Wajnberg, E., Fricaux, T., Taquet, N., Blomquist, G.J., Feyereisen, R., 2012. An insect-specific P450 oxidative decarbonylase for cuticular hydrocarbon biosynthesis. Proceedings of the National Academy of Sciences of the United States of America 109, 14858–14863.

Rewitz, K.F., O’Connor, M.B., Gilbert, L.I., 2007. Molecular evolution of the insect Halloween family of cytochrome P450s: phylogeny, gene organization and functional conservation. Insect Biochemistry and Molecular Biology 37, 741–753.

Riddick, D.S., Ding, X., Wolf, C.R., Porter, T.D., Pandey, A.V., Zhang, Q.Y., Gu, J., Finn, R.D., Ronseaux, S., McLaughlin, L.A., Henderson, C.J., Zou, L., Fluck, C.E., 2013. NADPH-cytochrome P450 oxidoreductase: roles in physiology, pharmacology, and toxicology. Drug Metabolism and Disposition 41, 12–23.

Romeis, J., Bartsch, D., Bigler, F., Candolfi, M.P., Gielkens, M.M., Hartley, S.E., Hellmich, R.L., Huesing, J.E., Jepson, P.C., Layton, R., Quemada, H., Raybould, A., Rose, R.I., Schiemann, J., Sears, M.K., Shelton, A.M., Sweet, J., Vaituzis, Z., Wolt, J.D., 2008. Assessment of risk of insect-resistant transgenic crops to nontarget arthropods. Nature Biotechnology 26, 203–208.

San Miguel, K., Scott, J.G., 2016. The next generation of insecticides: dsRNA is stable as a foliar-applied insecticide. Pest Management Science 72, 801–809.

Sanada-Morimura, S., Sakumoto, S., Ohtsu, R., Otuka, A., Huang, S.H., Thanh, D.V., Matsumura, M., 2011. Current status of insecticide resistance in the small brown planthopper, *Laodelphax striatellus*, in Japan, Taiwan, and Vietnam. Applied Entomology and Zoology 46, 65–73.

Sandstrom, P., Welch, W.H., Blomquist, G.J., Tittiger, C., 2006. Functional expression of a bark beetle cytochrome P450 that hydroxylates myrcene to ipsdienol. Insect Biochemistry and Molecular Biology 36, 835–845.

Sarapusit, S., Lertkiatmongkol, P., Duangkaew, P., Rongnoparut, P., 2013. Modeling of *Anopheles minimus* mosquito NADPH-cytochrome P450 oxidoreductase (CYPOR) and mutagenesis analysis. International Journal of Molecular Sciences 14, 1788–1801.

Sarapusit, S., Xia, C., Misra, I., Rongnoparut, P., Kim, J.J., 2008. NADPH-cytochrome P450 oxidoreductase from the mosquito *Anopheles minimus*: kinetic studies and the influence of Leu86 and Leu219 on cofactor binding and protein stability. Archives of Biochemistry and Biophysics 477, 53–59.

Schoville, S.D., Chen, Y.H., Andersson, M.N., Benoit, J.B., Bhandari, A., Bowsher, J.H., Brevik, K., Cappelle, K., Chen, M.M., Childers, A.K., Childers, C., Christiaens, O., Clements, J., Didion, E.M., Elpidina, E.N., Engsontia, P., Friedrich, M., Garcia-Robles, I., Gibbs, R.A., Goswami, C., Grapputo, A., Gruden, K., Grynberg, M., Henrissat, B., Jennings, E.C., Jones, J.W., Kalsi, M., Khan, S.A., Kumar, A., Li, F., Lombard, V., Ma, X., Martynov, A., Miller, N.J., Mitchell, R.F., Munoz-Torres, M., Muszewska, A., Oppert, B., Palli, S.R., Panfilio, K.A., Pauchet, Y., Perkin, L.C., Petek, M., Poelchau, M.F., Record, E., Rinehart, J.P., Robertson, H.M., Rosendale, A.J., Ruiz-Arroyo, V.M., Smagghe, G., Szendrei, Z., Thomas, G.W.C., Torson, A.S., Vargas Jentzsch, I.M., Weirauch, M.T., Yates, A.D., Yocum, G.D., Yoon, J.S., Richards, S., 2018. A model species for agricultural pest genomics: the genome of the Colorado potato beetle, *Leptinotarsa decemlineata* (Coleoptera: Chrysomelidae). Scientific Reports 8, 1931.

Schuler, M.A., 1996. The role of cytochrome P450 monooxygenases in plant-insect interactions. Plant Physiology 112, 1411–1419.

Shen, A.L., O’Leary, K.A., Kasper, C.B., 2002. Association of multiple developmental defects and embryonic lethality with loss of microsomal NADPH-cytochrome P450 oxidoreductase. The Journal of Biological Chemistry 277, 6536–6541.

Shi, L., Zhang, J., Shen, G., Xu, Z., Wei, P., Zhang, Y., Xu, Q., He, L., 2015. Silencing NADPH-cytochrome P450 reductase results in reduced acaricide resistance in *Tetranychus cinnabarinus* (Boisduval). Scientific Reports 5, 15581.

Snoeck, S., Kurlovs, A.H., Bajda, S., Feyereisen, R., Greenhalgh, R., Villacis-Perez, E., Kosterlitz, O., Dermauw, W., Clark, R.M., Van Leeuwen, T., 2019. High-resolution QTL mapping in Tetranychus urticae reveals acaricide-specific responses and common target-site resistance after selection by different METI-I acaricides. Insect biochemistry and molecular biology 110, 19–33.

Sun, X., Gong, Y., Ali, S., Hou, M., 2018. Mechanisms of resistance to thiamethoxam and dinotefuran compared to imidacloprid in the brown planthopper: Roles of cytochrome P450 monooxygenase and a P450 gene CYP6ER1. Pesticide biochemistry and physiology 150, 17–26.

Swevers, L., Huvenne, H., Menschaert, G., Kontogiannatos, D., Kourti, A., Pauchet, Y., ffrench-Constant, R., Smagghe, G., 2013. Colorado potato beetle (Coleoptera) gut transcriptome analysis: expression of RNA interference-related genes. Insect Molecular Biology 22, 668–684.

Tamura, K., Peterson, D., Peterson, N., Stecher, G., Nei, M., Kumar, S., 2011. MEGA5: Molecular evolutionary genetics analysis using maximum likelihood, evolutionary distance, and maximum parsimony methods. Molecular Biology and Evolution 28, 2731–2739.

Truman, J.W., Riddiford, L.M., 1999. The origins of insect metamorphosis. Nature 401, 447–452.

Velez, A.M., Fishilevich, E., Matz, N., Storer, N.P., Narva, K.E., Siegfried, B.D., 2016a. Parameters for successful parental RNAi as an insect pest management tool in Western corn rootworm, *Diabrotica virgifera virgifera*. Genes 8.

Velez, A.M., Jurzenski, J., Matz, N., Zhou, X., Wang, H., Ellis, M., Siegfried, B.D., 2016b. Developing an in vivo toxicity assay for RNAi risk assessment in honey bees, *Apis mellifera* L. Chemosphere 144, 1083–1090.

Wang, K., Peng, X., Zuo, Y., Li, Y., Chen, M., 2016. Molecular cloning, expression pattern and polymorphisms of NADPH-Cytochrome P450 reductase in the bird cherry-oat aphid *Rhopalosiphum padi* (L.). PloS One 11, e0154633.

Wang, M., Roberts, D.L., Paschke, R., Shea, T.M., Masters, B.S., Kim, J.J., 1997. Three-dimensional structure of NADPH-cytochrome P450 reductase: prototype for FMN- and FAD-containing enzymes. Proceedings of the National Academy of Sciences of the United States of America 94, 8411–8416.

Waskell, L., Kim, J.J.P., 2015. Electron transfer partners of cytochrome p450. in: Paul, R., Montellano, O.D. (Eds.). Cytochromes P450: Stucture, Mechanism and Biochemistry. Springer International Publishing, pp. 33–68.

Whyard, S., Singh, A.D., Wong, S., 2009. Ingested double-stranded RNAs can act as species-specific insecticides. Insect Biochemistry and Molecular Biology 39, 824–832.

Wybouw, N., Kosterlitz, O., Kurlovs, A.H., Bajda, S., Greenhalgh, R., Snoeck, S., Bui, H., Bryon, A., Dermauw, W., Van Leeuwen, T., Clark, R.M., 2019. Long-term population studies uncover the genome structure and genetic basis of xenobiotic and host plant adaptation in the herbivore *Tetranychus urticae*. Genetics 211, 1409–1427.

Zewen, L., Zhaojun, H., Yinchang, W., Lingchun, Z., Hongwei, Z., Chengjun, L., 2003. Selection for imidacloprid resistance in *Nilaparvata lugens*: cross-resistance patterns and possible mechanisms. Pest Management Science 59, 1355–1359.

Zhang, J., Khan, S.A., Hasse, C., Ruf, S., Heckel, D.G., Bock, R., 2015. Full crop protection from an insect pest by expression of long double-stranded RNAs in plastids. Science (New York, N.Y.) 347, 991–994.

Zhang, X., Wang, J., Liu, J., Li, Y., Liu, X., Wu, H., Ma, E., Zhang, J., 2017. Knockdown of NADPH-cytochrome P450 reductase increases the susceptibility to carbaryl in the migratory locust, *Locusta migratoria*. Chemosphere 188, 517–524.

Zhang, Y., Wang, Y., Wang, L., Yao, J., Guo, H., Fang, J., 2016. Knockdown of NADPH-cytochrome P450 reductase results in reduced resistance to buprofezin in the small brown planthopper, *Laodelphax striatellus* (fallen). Pesticide Biochemistry and Physiology 127, 21–27.

Zhao, G.Y., Liu, W., Brown, J.M., Knowles, C.O., 1995. Insecticide resistance in-field and laboratory strains of Western flower thrips (Thysanoptera, Thripidae). Journal of Economic Entomology 88, 1164–1170.

Zhao, J.Z., Bishop, B.A., Grafius, E.J., 2000. Inheritance and synergism of resistance to imidacloprid in the Colorado potato beetle (Coleoptera: Chrysomelidae). Journal of Economic Entomology 93, 1508–1514.

Zhu, F., Cui, Y., Walsh, D.B., Lavine, L.C., 2014. Application of RNAi towards insecticide resistance management. in: Chandrasekar, R., Tyagi, B.K., Gui, Z., Reeck, G.R. (Eds.). Short Views on Insect Biochemistry and Molecular Biology. Academic Publisher, Manhattan, USA, pp. 595–619.

Zhu, F., Moural, T.W., Nelson, D.R., Palli, S.R., 2016. A specialist herbivore pest adaptation to xenobiotics through up-regulation of multiple Cytochrome P450s. Scientific Reports 6, 20421.

Zhu, F., Sams, S., Moural, T., Haynes, K.F., Potter, M.F., Palli, S.R., 2012. RNA interference of NADPH-cytochrome P450 reductase results in reduced insecticide resistance in the bed bug, *Cimex lectularius*. PloS One 7, e31037.

Zhu, F., Xu, J., Palli, R., Ferguson, J., Palli, S.R., 2011. Ingested RNA interference for managing the populations of the Colorado potato beetle, *Leptinotarsa decemlineata*. Pest Management Science 67, 175–182.

Zhu, K.Y., Palli, S.R., 2020. Mechanisms, applications, and challenges of insect RNA interference. Annual Review of Entomology 65, 293–311.

Zotti, M., Dos Santos, E.A., Cagliari, D., Christiaens, O., Taning, C.N.T., Smagghe, G., 2018. RNA interference technology in crop protection against arthropod pests, pathogens and nematodes. Pest Management Science 74, 1239–1250.

